# Steric shielding of the KRAS4B hypervariable region enables isoform-specific inhibition of prenylation

**DOI:** 10.64898/2026.03.18.712636

**Authors:** Jonas N. Maskos, Yvonne Stark, Valentin L. Rohner, Andri Häfliger, Dorothea Winkelvoß, Kari Kopra, Peer R.E. Mittl, Andreas Plückthun

## Abstract

Mutant KRAS is a potent oncogene, serving as a tumor driver in many solid human cancers. Current small-molecule inhibitors target the highly conserved G-domain, but to gain further mechanistic insight into the roles of different isoforms, we investigated the strategy of sterically shielding the unstructured hypervariable regions (HVRs). KRAS HVRs undergo a series of post-translational modifications that enable intracellular trafficking and membrane attachment. Previous attempts to drug KRAS by preventing its post-translational modification, based on inhibition of the involved prenylation enzymes have been largely unsuccessful. In this study, we explored the property of Designed Armadillo Repeat Proteins (dArmRPs) to specifically bind unstructured regions. We assembled a dArmRP to recognize the unstructured KRAS4B-HVR and developed it into a high-affinity binder by directed evolution. The resulting dArmRP recognizes the 14 C-terminal residues of unprocessed KRAS4B, thereby blocking the farnesyltransferase-binding epitope. This steric shielding disrupts KRAS4B post-translational modification and thereby significantly reduces its plasma membrane localization, while demonstrating complete selectivity over KRAS4A, NRAS, and HRAS. This work establishes the shielding of intrinsically disordered regions as a precise biochemical strategy to control protein function and provides an isoform-specific tool to dissect KRAS biology.

**Graphical Abstract:** 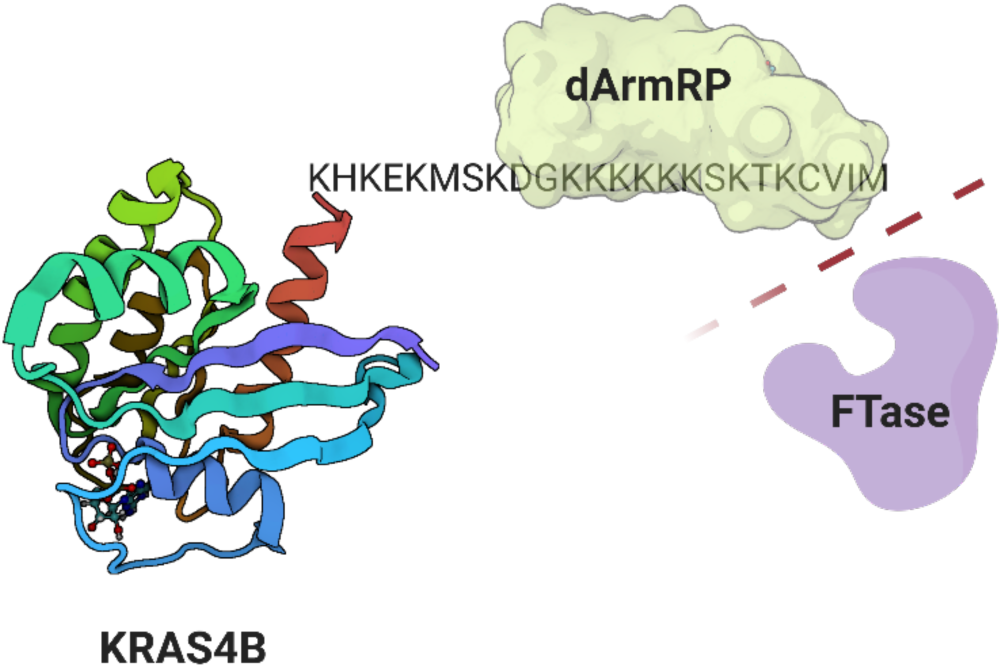

Graphical representation of how the unstructured KRAS4B-HVR is occupied by a dArmRP, making it inaccessible for the FTase.

## Introduction

The RAS family of small GTPases, encoded by the HRAS, NRAS, and KRAS genes, play central roles in transducing signals from membrane receptors to regulate cell growth and differentiation. Among these, KRAS is the most frequently mutated oncogene in human cancers. Alternative splicing of the KRAS pre-mRNA produces two isoforms, KRAS4A and KRAS4B, which share nearly identical catalytic G-domains but differ substantially in their hypervariable regions (HVRs) (1). The C-terminal HVRs are largely unstructured and contain membrane targeting sequences that are crucial for subcellular trafficking and localization. Both KRAS4A and KRAS4B are oncogenic when constitutively activated by mutations in exon 2 or 3, although they exhibit tissue-specific expression patterns and different functional characteristics in regulating downstream signaling pathways such as RAF-MAPK and PI3K (2). KRAS4B is a globular protein consisting of a structured GTPase domain (G-domain; residues 1-166) and the unstructured HVR (residues 167-188). The intrinsically disordered HVR, located at the C-terminus of KRAS4B, mediates anchoring to the plasma membrane. After translation, KRAS4B undergoes an extensive posttranslational modification process, during which the HVR is prenylated, proteolytically cleaved and carboxymethylated. In the initial rate-limiting step, the C-terminal CVIM motif is recognized by the cytosolic farnesyltransferase (FTase), and a farnesyl moiety is attached to the C185 residue via a thioether linkage (3, 4). In the absence of a functional FTase, KRAS4B can be alternatively geranylgeranylated by a cytosolic geranylgeranyl transferase (GGTase) (5). The prenylated KRAS4B then localizes to the endoplasmic reticulum, where the terminal VIM peptide is proteolytically cleaved off by the RAS converting enzyme 1 (RCE1), which specifically recognizes the prenylated CAAX motif (C =Cys, A=aliphatic [Leu, Ile, Val] and X=Met, Ser, Leu, Gln) (6). The prenylated and cleaved KRAS4B protein is recognized by an isoprenylcystein methyltransferase (ICMT), which methylates the carboxy group of the previously prenylated C185 (7). Finally, the chaperone phosphodiesterase 6δ (PDE6δ) recognizes the fully modified KRAS4B and directs it to the plasma membrane. Once at the membrane, the farnesyl moiety and the HVR’s hexa-lysine patch act together to secure stable anchorage (8, 9).

As KRAS requires prenylation for proper membrane localization and activation, the targeting of RAS modification and trafficking pathways has been investigated as a logical therapeutic approach. However, efforts to inhibit the initial prenylation step with single or combined inhibition of FTase and GGTase have largely failed in clinical trials against KRAS-driven tumors due to pathway redundancy and toxicity (10–12). Nevertheless, the FTase inhibitor tipifarnib has shown promising results in a phase II study for the treatment of HRAS mutant head-and-neck cancer and is being explored as a combination treatment with KRAS G12C inhibitors (13, 14). Pharmacological inhibition of the post-prenylation processing enzymes RCE1 and ICMT is only now being explored in preclinical studies due to a lack of potent, lead-like molecules and toxicity concerns (15). Additional preclinical approaches include the inhibition of trafficking chaperones, such as galectins and PDE6δ (16–18).

In this study, we explore the inhibition of KRAS4B prenylation by sterically shielding the HVR with synthetic binding proteins. To date, a whole set of synthetic binding proteins targeting different epitopes of the RAS G-domain has been developed that interfere with essential RAS functions such as nucleotide-exchange and downstream factor engagement (19). However, no approach for the inhibition of RAS membrane localization has been described. To address this gap, we exploit the unstructured nature of the HVR, utilizing the ability of designed armadillo repeat proteins (dArmRPs) to specifically recognize extended peptide stretches (20). In a related approach, de novo binders against the C-termini were designed, but their effect on prenylation was not investigated (21).

dArmRPs are a new class of engineered modular peptide binders based on the natural armadillo repeat protein scaffold such as importin-α and β-catenin (22). They consist of tandem armadillo repeats that form a superhelical structure with a peptide-binding groove. By stacking multiple repeats together with an N- and C-terminal capping repeat, an elongated binding surface is formed that can recognize extended peptide stretches in a modular manner. The modularity allows generating sequence-specific binders by combining modules selected for different target amino acid sidechains. The peptide binds in an antiparallel arrangement to the dArmRP.

A module consists of two dArmRP repeats that create a binding interface for two amino acid side chains of the target peptide (23). The consensus-derived dArmRP scaffold recognizes alternating arginine and lysine residues in an upper pocket (binding arginine in importin-α) and a lower pocket (binding lysine in importin-α), in a highly modular fashion (24). Furthermore, by utilizing a selection-based approach, new modules can be generated to recognize individual amino acids (25). By combining these specific modules, sequence-specific binders can theoretically be assembled for any extended peptide sequence, provided the necessary preexisting modules are available.

Here, we report for the first time the design and affinity maturation of a specific dArmRP to a natural target, KRAS4B-HVR, and assess its potential to interfere with KRAS4B prenylation and membrane attachment, providing a novel approach for isoform-specific targeting of this key oncogene.

## Results

### Initial design of a binder to the HVR of KRAS4B

We hypothesized that the unstructured nature and highly positive charge of the KRAS4B-HVR could serve as an ideal target for dArmRPs to sterically shield the region from normal cellular processing.

Previously, we described consensus dArmRPs with up to six internal modules that recognize extended peptide stretches composed of alternating arginine and lysine residues, while also tolerating arginine to lysine substitutions with only slightly compromised affinity (23, 24). In addition, we recently developed modules with high specificity and affinity for serine and threonine by yeast display (Stark et al., unpublished data). Based on these available design elements, we assembled an initial candidate, termed KB2, intended to recognize the 12 C-terminal residues (172–183) of the KRAS4B-HVR. To achieve this, we combined three Ser/Thr-binding modules with three Arg/Lys-binding consensus modules, resulting in a dArmRP with six internal modules (Figure 1A).

**Figure 1.**
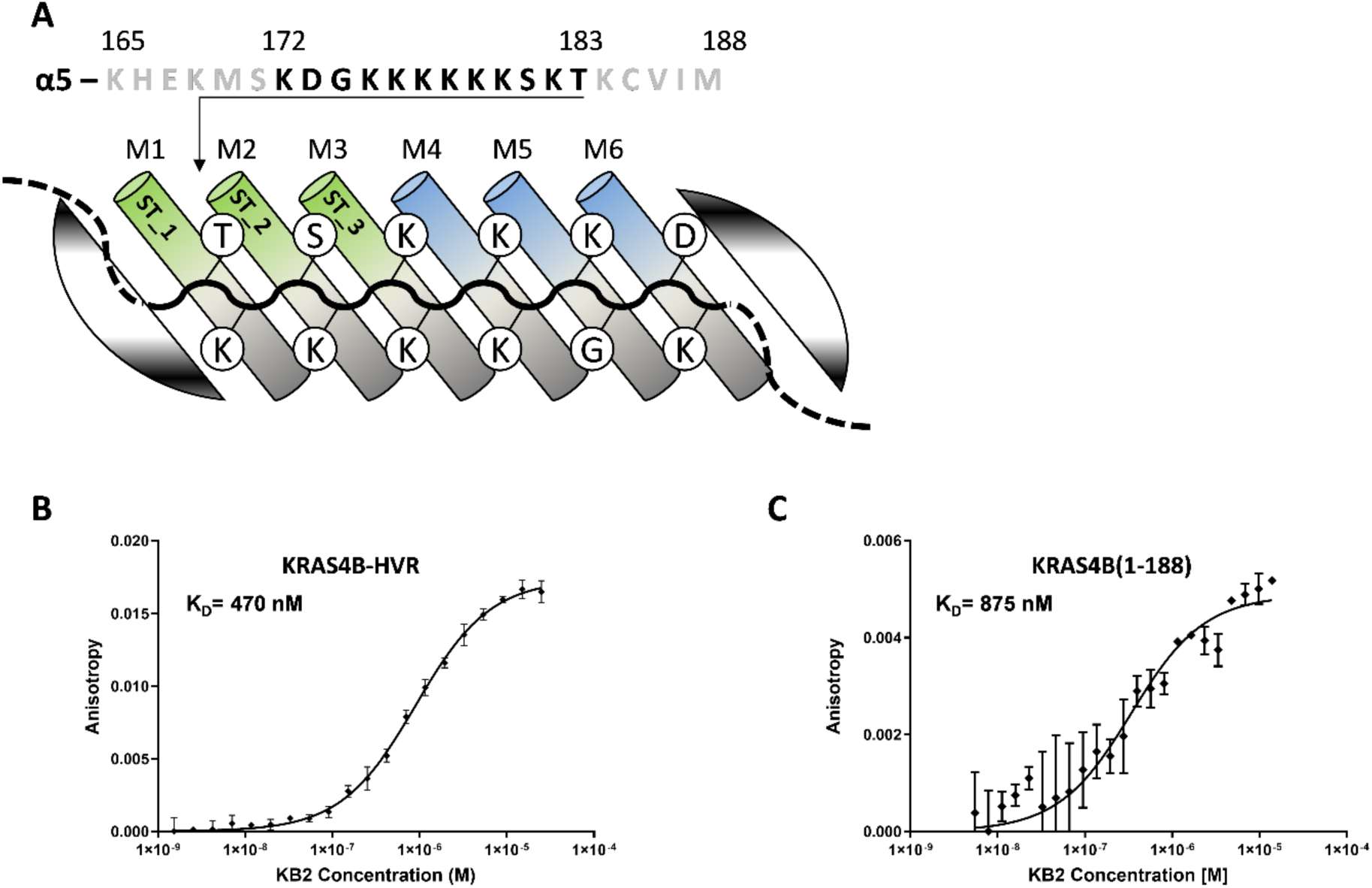
Assembly of a KRAS4B-HVR binding dArmRP. **(A)** Antiparallel binding of the KB2 dArmRP to residues 172-183 of the KRAS4B-HVR. The six structural modules of the dArmRP are shown as cylinders. Green modules represent selected modules for the binding of serine and threonine residues, while blue ones represent consensus modules. The resulting pockets between two structural modules are divided into the upper (arginine) and the lower (lysine) pocket. The N- and C-terminal capping modules are illustrated as striped areas. The peptide backbone bound on the dArmRP is indicated as a solid line. The dashed line indicates unbound residues exiting the binding interface, shown in the above sequence in grey. **(B and C)** Affinities of KB2 for KRAS4B (1-188) and the KRAS4B-HVR (165-188) determined by fluorescence anisotropy. The lower maximal anisotropy amplitude in (C) reflects the larger initial size of the full-length KRAS4B reporter protein, resulting in a smaller relative change in tumbling dynamics upon complex formation.

Notably, creating two distinct pockets for the binding of serine and threonine required careful structural integration of the first three modules. Because the binding surface for a single target residue is formed by two neighboring modules, positions 29 and 33 within the internal repeats participate in binding two adjacent target residues simultaneously; we term these “shared residues”. Therefore, the exact pocket selected for Ser/Thr binding was directly used for the modules one (ST_1) and two (ST_2), while module three (ST_3) was designed as chimera between the newly selected threonine pocket and the consensus Arg/Lys pocket. The designated KRAS4B-HVR target sequence also contains a glycine and an aspartic acid residue, for which no modules were available at the time of design. We assembled KB2 using the consensus module at these positions, anticipating that, while the scaffold might tolerate these residues without severe steric repulsion at the edge of the interface, achieving high-affinity target engagement would ultimately require directed evolution.

To validate the design, we evaluated the binding of KB2 to the peptide sequence of the KRAS4B-HVR(165–188) and the full-length KRAS4B protein with a K_D_ of 470 nM and 875 nM, respectively (Figure 1B,C). The affinity for both was measured by fluorescence anisotropy (FA), using either EGFP or mNeonGreen as N-terminal fusions to KRAS(1–188) and the KRAS4B-HVR.

### Affinity maturation of a KRAS4B-HVR binding dArmRP

While we were able to generate a binder to the KRASB-HVR by rational design, its affinity was clearly not high enough to compete with farnesyl- and geranylgeranyl transferases, which exhibit single-digit nanomolar affinities for KRAS4B (26, 27). We therefore set out to improve the binding affinity of KB2 in a directed evolution approach. A randomized library of the KB2 dArmRP was generated by error-prone PCR and used to select improved variants by yeast-surface-display (YSD) selection (25). In addition to the emergence of mutations, the sequence homology of the individual modules allowed for module shuffling as well as the extension of the initial construct by additional modules during the generation of the library (Figure 2A).

**Figure 2.**
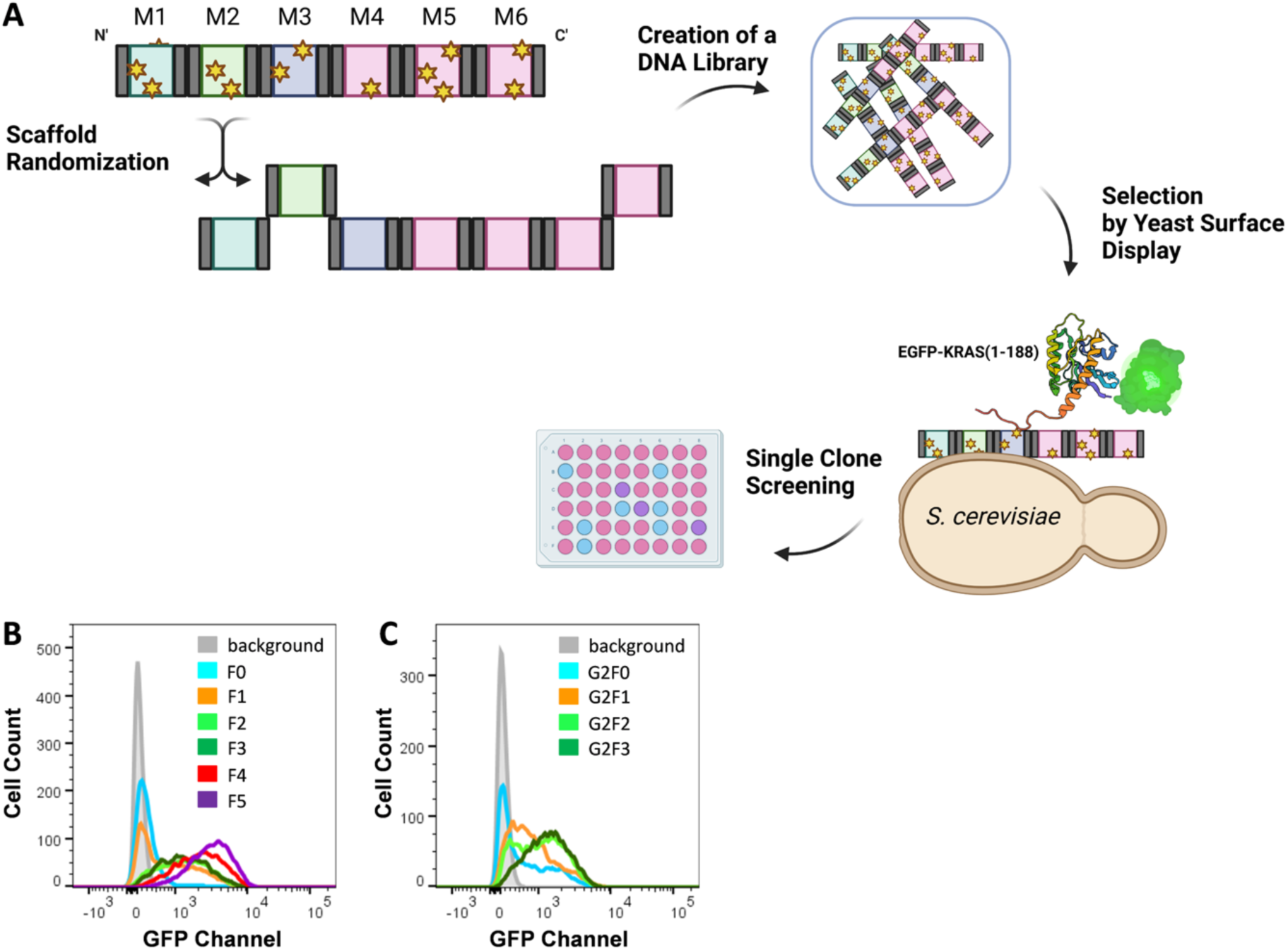
Affinity maturation of a KRAS4B-HVR binding dArmRP. **(A)** A DNA library was generated by randomization of the KB2 dArmRP by the introduction of mutations by error-prone PCR (yellow stars), and repeat shuffling. The homologous regions of the modules are indicated by grey vertical bars. Blue, green and turquoise shaded blocks symbolize repeats derived from serine/threonine-binding modules, and pink shaded blocks symbolize consensus repeats. dArmRPs with a strong binding to EGFP-KRAS (1-188) are enriched by YSD and FACS**. (B)** Enrichment of target-binding yeast cells over the first five rounds of YSD at 500 nM EGFP-KRAS4B. **(C)** Enrichment of target-binding yeast cells over three additional rounds of YSD after the 2^nd^ scaffold randomization 31 nM EGFP-KRAS4B.

dArmRP variants were selected for their binding to GTPγS-loaded EGFP-KRAS4B(1–188) in five initial rounds before the selected pool was re-randomized and subjected to three additional rounds of selection. To enforce specific binding to KRAS4B-HVR, we included a non-fluorescent sfGFP(Y66A) fused to a related, highly-positively charged peptide and cell lysate of KRAS-deficient HeLa cells as competitors in the selection. All details on the selection conditions are shown in Table S1. We observed a continuous improvement in binding signals of the selected pools after both randomization steps, allowing a reduction of target concentrations from 750 to 31 nM during the selection process (Figure 2B-C). To identify the most affine binders from the selected pool, we performed single-clone analysis of the yeast-displayed clones by flow cytometry before characterizing the top clones as recombinant proteins expressed in *E. coli*.

We isolated KRAS4B-binding dArmRPs with six to eight internal modules that exhibited two major changes when compared to the original KB2 design. First, we found a dominant S25R mutation in the N-terminal capping repeat of the selected dArmRPs present in >98 % of all sequences (see structural interpretation further below). Further, we observed a shuffling of the modules. In all selected clones, the last three C-terminal modules are consensus modules, while the N-terminal modules are derived from the serine/threonine-binding modules. Interestingly, the first module ST_1 was strongly deselected and found to be absent in all selected dArmRPs. Instead, the selected dArmRPs contain up to five ST_3 modules or alternating ST_3 and ST_2 modules. The individual dArmRPs did acquire additional point mutations without apparent pattern or evident effect on the function.

### Evolved dArmRPs selectively bind unmodified KRAS4B

The selected and evolved KRAS4B-binding dArmRPs were further subjected to in-depth characterization. Specific binding to the unmodified KRASB-HVR(165–188) was confirmed by performing enzyme-linked immunosorbent assays (ELISA) with KRAS4B(1–188), HVR-truncated KRAS4B (1–166) and fully-modified KRAS4B(1-185 FMe). Binding was only observed for KRAS4B(1–188) (Figure 3A). To further confirm binding selectivity, we expressed KRAS and the isoforms NRAS and HRAS in HEK293T cells produced in HEK293T cells (the different RAS HVRs are shown in Figure S1). We then probed the resulting cell lysates by western blot, both in the presence and absence of the dual FTase/GGTase inhibitor FGTI-2734 (28). The selected dArmRPs clearly recognize overexpressed KRAS4B when its prenylation is pharmacologically suppressed, while no cross-reactivity for the isoforms NRAS and HRAS can be observed under the same condition. Similarly, no binding for regularly expressed — and thus modified — KRAS4B can be observed (Figure 3B, C). These results confirm that the selected dArmRPs specifically recognize the unmodified C-terminus of KRAS4B.

**Figure 3.**
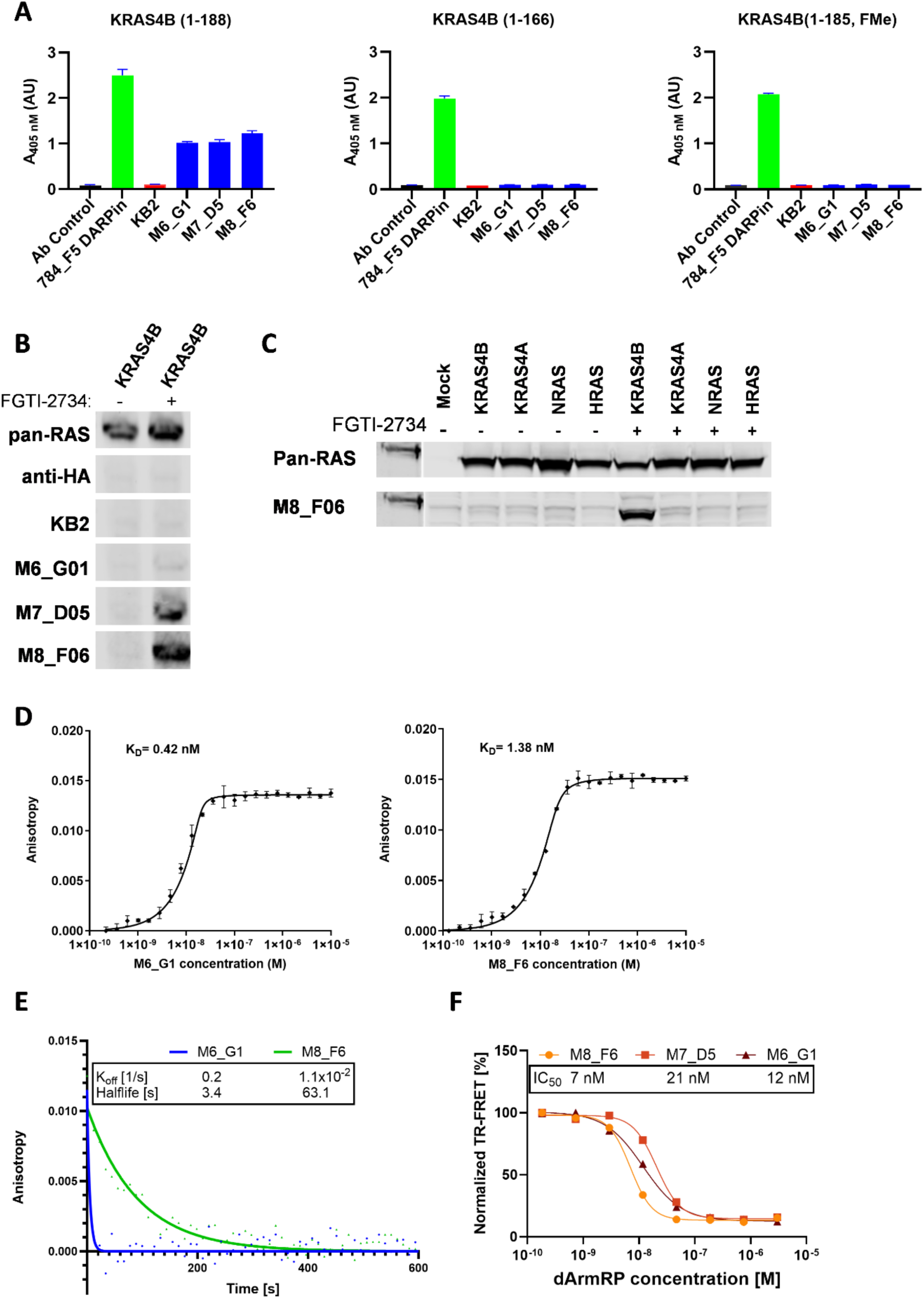
Biochemical characterization of the selected and evolved dArmRPs. **(A)** Binding of dArmRPs with six to eight internal repeats to KRAS4B(1-188), KRAS4B(1-166) and KRAS4B(1-185, FMe) in ELISA. The DARPin 784_F5, binding to the RAS G-domain, represents a positive control. Ab control denotes the secondary antibody only. **(B)** Detection of overexpressed KRAS4B after treatment with the prenylation inhibitor FGTI-2734. HA-tagged dArmRPs were used as primary detection reagents. A pan-RAS antibody serves as positive control. **(C)** Detection of overexpressed RAS isoforms by M8_F06-AzDye680. **(D)** Affinities of the dArmRPs M6_G1, and M8_F6 for KRAS4B-HVR(165-188), determined by FA. **(E)** Dissociation rates of the of the dArmRPs M6_G1, and M8_F6, for KRAS4B-HVR, determined by FA. **(F)** Inhibition of SOS-mediated nucleotide exchange by the dArmRPs M6_G1, M7_D5 and M8_F6.

We further determined the affinities of the selected dArmRPs by FA to estimate whether they would be able to compete with the prenylating enzymes for the KRAS4B-HVR. All characterized binders exhibited a K_D_ between 0.4 and 9 nM for the KRAS4B-HVR, providing a 100- to 1000-fold increase in affinity compared to the parental binder KB2 (Figure 3D, Table S2). Interestingly, the observed equilibrium affinities did not vary significantly between dArmRPs with different numbers of internal modules, even though the binding signals observed in ELISA and western blots strongly correlated with the number of internal modules. We could subsequently show that dArmRPs with more internal repeats do exhibit a slower k_off_ despite a similar global K_D_ (Figure 3E), suggesting that the dissociation rate is the key quantity determining sensitivity in western blots.

To assess whether the binding of its HVR interferes with essential functions of KRAS4B we measured SOS-mediated nucleotide exchange in the presence of the selected dArmRPs. Interestingly, we found that these synthetic binding proteins are potent inhibitors of nucleotide exchange with IC_50_ values of 7 to 21 nM (Figure 3F). This observation was unexpected, since the HVR as an extension of the α5 helix is rather distant from the SOS1/SOS2-KRAS binding interface, which is the switch I/II region.

### Structural basis of the dArmRP interaction with the KRAS4B-HVR

To determine how the evolved dArmRPs recognize the KRAS4B-HVR, we explored the structures of the complexes using both X-ray crystallography and AlphaFold3 (AF3) modeling (29). As it proved challenging to obtain well-diffracting protein crystals due to the high solubility of standard dArmRPs (30) we engineered a specific variant to overcome this limitation. We introduced two charge-reducing mutations per module (E2K, D9N) on the “backside” of the protein, away from the binding interface. We thereby intended to reduce its high solubility, and we replaced the N-terminal cap with a stabilized variant to enhance its rigidity (31). This modified backbone yielded two crystal forms of a six-module dArmRP (M6_G1) bound to a synthetic KRAS4B peptide (residues 175-188), diffracting to 2.07 and 2.85 Å resolution, respectively (Figure 4A, Table S3). Interestingly, we observed near-atomic accuracy when comparing the experimental 2.07 Å resolution structure to an AF3-derived model of the complex, with a RMSD of 0.8 to 1.0 Å for the dArmRP-peptide complex and a RMSD of 0.7 to 1.2 Å for the peptide only (Figure 4B). Notably, a similar extent of variation is found for the structures of the six dArmRP-peptide complexes within the crystal’s unit cell. This structural agreement validated our use of AF3 to predict the structures of the other evolved variants.

**Figure 4.**
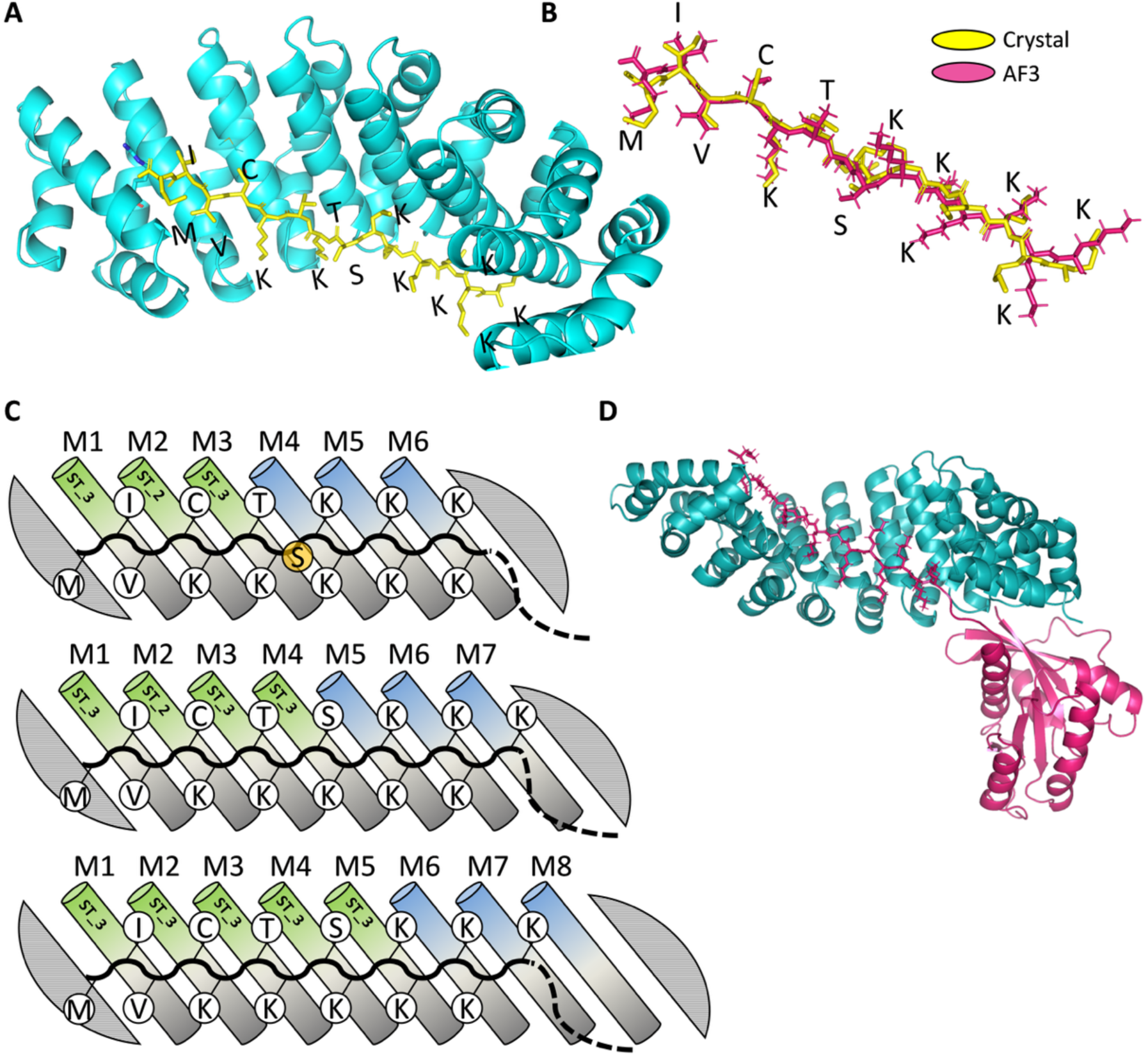
Binding mode of selected dArmRPs to the KRAS4B-HVR. **(A)** 2.07 Å resolution **c**rystal structure of the modified M6_G1 dArmRP (cyan) bound to a peptide comprised of residues 175-188 of KRAS4B (yellow). **(B)** Overlay of the bound HVR peptide obtained from the crystal structure (yellow) and from a AF3-generated model (pink). **(C)** Schematic illustration of the dArmRP-peptide interaction for the binders M6_G1, M7_D5 and M8_F6, from top to bottom, as deduced from AF3. The unbound S181 is highlighted in orange for the dArmRP with six repeats. **(D)** AF3-generated model of M8_F6 (cyan) bound to KRAS4B (pink).

The combination of experimental structure determination and computational modeling revealed that affinity maturation induced a register shift of the bound KRAS4B-HVR peptide, rather than exclusively optimizing localized side-chain contacts. In contrast to our initial KB2 design, the evolved dArmRPs shifted the binding register by four amino acids. The 2.07 Å structure demonstrates that the C-terminal M188 acts as the first engaged amino acid relative to the dArmRP N-terminus, with the bound sequence extending to K175. This C-terminal stretch of 14 residues is consistently observed or predicted as the binding epitope for all analyzed dArmRPs, independent of the number of internal modules (Figure 4C).

This register shift in the complex structure is anchored by a dominant S25R mutation in the N-terminal capping repeat, a substitution present in over 98% of all enriched sequences. Our structural analysis provides the molecular basis for this selection enrichment: the acquired arginine residue forms a salt bridge with the free carboxyl group of the C-terminal M188, allowing it to fold onto helix three of the N-terminal cap (Figure S2A). Furthermore, the shuffling and multiplication of the ST_3 module during library generation accommodated this new register. Originally incorporated to recognize T183 and S181, the duplicated ST_3 modules in the shifted frame provide an extended complementary surface that enables binding of I175 and C185.

A comparison of the predicted structures of variants with six to eight internal repeats highlights the structural differences due to binding surface length. Variants with six repeats sacrifice S181 binding to accommodate an extra lysine, while seven- and eight-repeat variants engage all 14 residues. The eighth repeat in particular allows K175 to bind in the consensus arginine pocket rather than between the seventh repeat and the C-terminal cap. Predictions for the complex of selected dArmRPs with either the KRAS4B-HVR peptide (174–188) or the whole KRAS4B protein do not differ in the placement of the HVR on the dArmRP surface (Figure 4D, S2B-C). The KRAS4B-HVR exits the dArmRP binding surface, starting with G173 at the level of the last internal module, beginning to adopt a helical structure from residues D172-M170. The predicted structures suggest a high degree of freedom for the relative position of the KRAS4B G-domain to the dArmRP.

Finally, we used these structural insights to evaluate the potential for off-target binding. A protein BLAST search using the identified 14-residue target sequence (KRAS4B 174-188) returned high homology primarily for the extended polybasic region (residues 175-185) of other cellular proteins, but critically, these sequences lacked homology to the C-terminal CVIM motif (Figure S3). Because our biochemical data demonstrated that even the absence of the three C-terminal residues of the KRAS4B results in a 60-fold loss in binding affinity, it can be considered unlikely that the selected dArmRPs exhibit significant off-target binding to other cellular proteins.

### Selected dArmRPs prevent post-translational modification and membrane trafficking of KRAS4B

To explore the potential of the selected KRAS4B-binding dArmRPs to interfere with KRAS4B processing *in cellulo*, we utilized microinjection coupled with live-cell microscopy (Figure 5A, B). First, to demonstrate prevention of KRAS4B-processing, EGFP-fused KRAS4B-HVR(165–188) and mCherry-fused dArmRP were microinjected in HEK293 cells. This allowed precise control over the relative abundancies of both proteins in the cell as well as the monitoring of KRAS4B processing based on the ratio of cytosolic and membrane-localized KRAS4B-HVR. When injected alone, the fluorescent HVR localized almost exclusively to the plasma membrane within 1.5 hours, indicating efficient post-translational processing by the endogenous cellular machinery (Figure 5A). Strikingly, when co-injected with a three-fold excess of KB_M8_F6, the HVR was entirely retained in the cytosol, with no membrane localization observed over a 12-hour period (Figure 5C).

**Figure 5.**
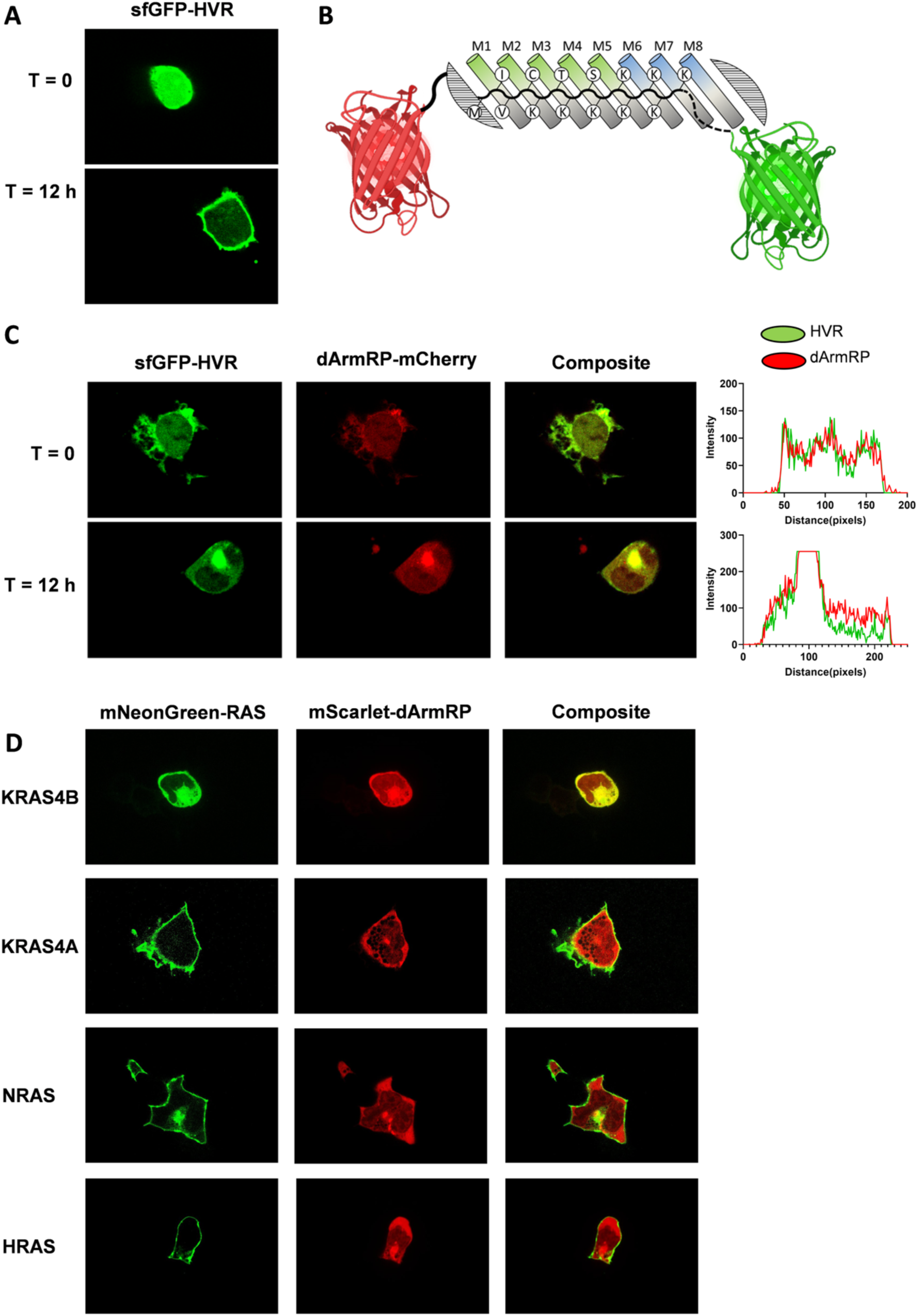
Inhibition of KRAS4B prenylation by selected dArmRPs. **(A)** Localization of recombinantly expressed sfGFP-HVR(165-188) directly and 12 h after microinjected into HEK293T cells. **(B)** Schematic representation of the experimental setup. In all shown experiments the RAS isoforms are fused to red fluorescent protein (mCherry, mScarlet) and the tested dArmRP are fused to a green fluorescent protein (sfGFP, mNeonGreen) which should results in co-localization of both colors upon complex formation. **(C)** Localization of recombinant sfGFP-HVR (green) directly and 12 h after microinjected as a protein into HEK293T cells in complex with mCherry fused dArmRP (red) M8_F6. The colocalization of both proteins is shown as an overlay of both channels and quantified along a line drawn across the displayed cell. **(D)** Localization of mNeongreen-fused RAS isoforms or mScarlet-fused dArmRP KB_M8_F6 12 h after microinjection of the respective plasmids.

While this protein co-injection confirmed that the dArmRP efficiently shields the isolated HVR, we sought to validate this effect in the context of the full-length KRAS4B protein. Interestingly, we discovered that microinjected recombinant full-length KRAS4B fails to reach the plasma membrane, whereas KRAS4B translated de novo from a microinjected plasmid traffics properly (Figure S4). This observation suggests that the required posttranslational modifications must occur early after translation, and perhaps in a co-translational manner, which to date has not been reported.

Consequently, to evaluate our binder in a native translational context and to confirm isoform specificity, we adapted our assay to a plasmid co-expression model. We microinjected pcDNA 3.1 plasmids encoding mNeonGreen-fused full-length RAS isoforms alongside mScarlet-fused dArmRP and monitored RAS protein localization (Figure 5D). As expected, dArmRP M8_F6 strongly co-localized with KRAS4B, sequestering a substantial fraction of the oncoprotein in the cytosol. Crucially, this cytosolic sequestration was completely absent for the highly homologous isoforms KRAS4A, NRAS, and HRAS, demonstrating exquisite isoform selectivity. We note that in this steady-state co-expression setup, a subset of KRAS4B still achieved plasma membrane association, highlighting the kinetic challenge of fully outcompeting endogenous processing enzymes during *de novo* translation. Taken together, these experiments clearly demonstrate the ability of selected dArmRP to selectively block KRAS4B processing in a cellular context while sparing all other RAS isoforms.

## Discussion

In this study, we describe the assembly and affinity maturation of a dArmRP for the binding of the KRAS4B-HVR with the aim to shield it from prenylating enzymes. For the dArmRP assembly, available modules compatible with the polar amino acids threonine and serine were combined with the consensus module that mediates binding to arginine and lysine residues. As the assembled dArmRP exhibited only high-nanomolar affinity, we performed affinity maturation and obtained single-digit nanomolar binders (Figure 2, 3D). In general, an affinity maturation step is not intended in assembling a dArmRP but was required due the limited availability of modules for the amino acids to be bound at the time the study was initiated. The subsequent analysis then showed that the increase and rearrangement of modules was a direct consequence of the selection pressure for affinity and slower dissociation, and the structural analysis explained why it was a solution with better complementarity to the sequence to be bound than any point mutation that would have been possible.

Crystallographic studies and AF-generated models revealed that the selected dArmRP recognize and are complementary to the 14 residues 175-188 of the KRAS4B-HVR (Figure 4A, B). In contrast to the initial design, the binding register shifted by four amino acids, with the residue I187 occupying the first pocket. The binding of the additional residues results from the shuffling and multiplication of the first three modules, initially selected for the binding of serine and threonine, creating additional compatibilities. Thereby, the additional modules allow an optimal placement of all 14 residues on the dArmRP’s binding surface (Figure 4C). Further, the acquired S25R mutation in the N-terminal cap provides a salt-bridge to the carboxy-terminus of the HVR, as an additional interaction (Figure S2A). The changes incorporated during the affinity maturation illustrate how the interaction surface between the dArmRP and the HVR was maximized for a gain in affinity.

We show that the selected dArmRPs discriminate unmodified KRAS4B(1–188) from fully modified KRAS4B(1-185, FMe) (Figure 3A). They can thus be utilized as a detection reagent for unmodified KRAS4B when sufficiently abundant, as they do not recognize the modified form (Figure 3B). The specific binding of the unmodified KRAS4B-HVR(165–188) consequently translates into prevention of its prenylation as shown in cellular studies. As the outcome in this system is obviously determined by the relative abundancies of KRAS4B and dArmRP we additionally showed that the most potent dArmRP, M8_F6, does completely block KRAS4B membrane association over 12 h when microinjected as a complex (Figure 5C). The described functional assays highlight the phenotypic differences between the dArmRPs with six to eight modules. Despite their similar equilibrium K_D_, the dArmRP M8_F6 shows a 20-fold decrease in k_off_ over that of M6_G1 (Figure 3E), which results in superior performance in assays that involve washing or competition.

Interestingly, we observed that binding of the KRAS4B-HVR by the selected dArmRPs leads to potent inhibition of SOS-mediated nucleotide-exchange (Figure 3F). This finding was unexpected based on the distance of both binding epitopes, the large size of the SOS1/2 proteins might still lead to steric clashes with the bound dArmRP as seen in Figure S2D. Alternatively, the rigid fixation of the HVR in an extended conformation distant from the G-domain might allosterically alter the dynamics in the KRAS4B switch regions, thereby affecting SOS engagement. It can only be speculated whether the inhibition of nucleotide-loading of KRAS4B by dArmRP binding upon KRAS4B translation could result in a phenotypic effect. However, the reported instability of nucleotide-free RAS could potentially promote its degradation (32).

Translating this potent *in vitro* profile into complete phenotypic suppression of KRAS4B activity *in cellulo*, however, remains a kinetic challenge. The quantitative shielding of a substrate (the KRAS4B-HVR) from its modifying enzymes (FTase and GGTase) requires the binder to completely outcompete an irreversible enzymatic modification. Our microinjection experiments demonstrate that when high local concentrations and favorable stoichiometries are achieved, dArmRP M8_F6 can successfully and selectively block KRAS4B membrane association for up to 12 hours. Yet, achieving this blockade continuously in a steady-state cellular environment is limited by the binder’s dissociation rate. While the dArmRP M8_F6 exhibits an improved k_off_ (1.1x10^-2^ s^-1^) over variants with fewer modules (Figure 3E), this dissociation rate is still relatively fast compared to typical values for antibodies or designed ankyrin repeat proteins (DARPins) binding to globular proteins, which often exhibit dissociation rates in the range of 10^-3^ to 1 x 10^-5^ s^-1^ also in the context of RAS binding (33, 34). Future efforts towards improvement of the dArmRP’s dissociation rate would likely result in more potent inhibition of KRAS4B prenylation.

Taken together, we report the engineering of a (splice)-isoform-specific affinity reagent for KRAS4B that inhibits its prenylation by sterically shielding the KRAS4B-HVR from prenylating enzymes, providing a novel mechanism for targeting of the KRAS oncoprotein. Its potential therapeutic use is currently limited by the lack of technologies that enable efficient cytosolic delivery of such protein drugs and the relatively fast dissociation rate of the selected dArmRPs. Nonetheless, a variety of viral and non-viral delivery systems are currently being developed (35, 36) that could be co-opted for this purpose. As an additional strategy, we propose that retargeting of E3-ligases (37) or RAS-specific proteases (38) for the degradation of KRAS4B could be achieved with the selected KRAS4B-specific dArmRPs to augment phenotypical effects. Especially in the context of enzyme retargeting, the dArmRP’s fast dissociation rate could be beneficial, enabling a high turnover rate. In this enhanced format, the selected dArmRPs could serve as valuable tools to study the roles of the splice-isoforms KRAS4B/4A in the context of KRAS-mutant cancers which, despite increasing attention, is still understudied (2).

## Materials and Methods

### Molecular cloning

The DNA for the initial sequence of KB2 was ordered at Twist Bioscience and cloned into the in-house plasmids pQiq-HA and pYD2 using BamHI and HindIII restriction sites. The vector pQiq-HA is a derivative of the vector pQE30 (Qiagen) (39) with a 3C-protease cleavable N-terminal MRGSHis_6_- and a C-terminal HA-tag, carrying an ampicillin resistance. The vector pYD2 is a derivative of the vector pCTCON2 (40) with additional BamHI and HindIII restriction sites directly at the N- and C-termini of the protein of interest and optimized flanking sequences for homologous recombination of dArmRPs.

After the selection by yeast surface display (see below), screening hits were purified from the respective yeast cell cultures using the Zymoprep^TM^ Yeast Plasmid Miniprep I Kit (Zymo Research). The pYD2_dArmRP plasmids were used to transform *E. coli* XL1-Blue cells for amplification, and dArmRPs were subsequently subcloned into the plasmid pQiq-HA via BamHI and HindIII restriction sites.

### Protein expression and purification

All proteins were expressed in *E. coli* BL21-Gold(DE3) cells and all expression constructs are mentioned in Table S4. dArmRPs and peptide-sfGFP or -mCherry fusions were expressed as previously described (41). EGFP-KRAS(1–188) was expressed for 18 h at 18°C in Terrific Broth containing 2% (v/v) EtOH. Expression of KRAS was induced with 0.2 mM IPTG after the culture had reached OD_600_ = 0.8. All steps of protein purification were performed at 4°C. Bacterial pellets were resuspended in lysis buffer (50 mM Tris (pH 8.0), 10% (w/v) glycerol, 300 mM NaCl, 20 mM imidazole; 10 mM MgCl_2_, 0.1 mM GDP containing 25 ng/ml DNAse, 1 µg/ml lysozyme, 5 µg/ml leupeptin, 48 µg/ml Petabloc SC, and 1 µg/ml pepstatin A) and lysed via sonication. After centrifugation (20,000 × g, 30 min) and filtration (0.2 µm), KRAS was purified using Ni-NTA metal-ion affinity chromatography (IMAC). Subsequently, the IMAC eluate was dialyzed against 50 mM HEPES (pH 7.4), 10% (w/v) glycerol, 2 mM TCEP, 25 mM NaCl, 20 mM imidazole; 10 mM MgCl_2_ involving cleavage of the N-terminal tag by tobacco etch virus (TEV) protease. TEV protease and uncleaved protein were removed by reverse IMAC. KRAS was further purified by cation exchange (Mono S 5/50 GL) and size exclusion chromatography (HiLoad 16/600 Superdex 75 pg) to provide a full-length, monomeric protein. All proteins were analyzed by ESI-MS, confirming the expected molecular weight.

### Library generation

For this project two libraries were generated, an initial library (G1 F0) and a second generation library (G2 F0), where the G2 F0 library was created after five selection rounds on the initial library. Both the initial library as well as the re-randomized library were generated by error-prone PCR (EP-PCR) using the GeneMorph II Random Mutagenesis Kit (Agilent). The EP-PCR was performed on the whole coding sequence of either the parental sequence of the initially designed dArmRP or the pool of dArmRP sequences present after sorting round five. EP-PCR parameters were chosen to yield 1-2 mutations per sequence for the initial library and 4-5 mutations for the re-randomization according to the manufacturer’s protocol (Table S5). The primers used include flanking regions of 144 bps upstream and 151 bps downstream of the dArmRP coding region, which enable yeast-mediated homologous recombination into the yeast-surface-display vector (pYD2).

Randomized dArmRP fragments generated by the EP-PCR were used to transform *S. cerevisiae* EBY100 cells together with linearized (XhoI, PstI) pYD2_spacer plasmid by square wave electroporation as described previously (25).

### Yeast surface display (YSD)

YSD was essentially performed as described previously (25). Briefly, dArmRP proteins are C-terminally fused to Aga2p. Flanking the dArmRP, an N-terminal HA tag and a C-terminal c-myc tag are present that can be detected by the respective anti-tag antibodies, with the secondary antibody being fluorescently labeled for flow cytometry detection. A yeast surface expression control is performed for all native libraries generated, by parallel detection of the HA and c-myc tag. During selections, only the C-terminal c-myc tag is detected to allow gating for cells expressing a full-length construct along with the target signal. Analytical flow cytometry measurements were performed on a FACSCanto II cytometer (BD Biosciences) and FACS was performed on a FACSAria^TM^ III sorter (BD Biosciences). Flow cytometry and FACS data were analyzed with FlowJo (V10).

Five rounds of YSD sorting (1.1-1.5) were performed on the initial library (G1 F0) with stepwise decreasing target protein concentrations of 750 to 62.5 nM using a FACSAria^TM^ III sorter (BD Biosciences) followed by three additional rounds (2.1-2.3) on the second-generation library (G2 F0) with target concentrations being reduced from 62.5 to 31.25 nM. Competition with the peptide (KA)_3_KRKLSF fused to darkGFP (42) in 10-20-fold excess over the target protein was included in rounds 1.3, 2.2, and 2.4 to eliminate dArmRPs that might recognize GFP. Additionally, cell lysate derived from HeLa cells with a KRAS knockout (Abcam) was added in rounds 1.2-1.4 and 2.3 to deplete dArmRPs that recognize frequently expressed human proteins or that lose target-binding in their presence. The detailed conditions of each round are described in Table S1.

To isolate promising candidates, the pool obtained from the last sorting round was plated and 96 colonies were picked for single clone analysis by flow cytometry. The binders giving the highest target signal were subcloned into a bacterial expression vector and expressed in a small-scale format.

### Fluorescence anisotropy

Selected dArmRPs were characterized by measuring affinities by fluorescence anisotropy (FA). The assay was performed in black non-binding 96-well plates (Greiner bio-One) in a final volume of 100 μl. All measurements were performed in PBS (137 mM NaCl, 3 mM KCl, 8 mM Na_2_HPO_4_, 1.5 mM KH_2_PO_4_), supplied with 0.05 % (v/v) Tween-20. A dilution series of 24 different dArmRP concentrations was measured against a constant concentration of peptide-mNeongreen or -GFP fusion. The reporter peptide concentration was chosen to be 40 nM. Starting concentrations of the dArmRP dilution series ranged from 10 to 25 μM, 0.6-fold dilution steps were used to titrate the dArmRPs. Each measurement was performed in duplicates. The anisotropy was measured on a Tecan Spark reader (Tecan) equipped with a fluorescence polarization module. The duplicate measurements were averaged and normalized by subtraction of the lowest anisotropy value from all other values. A 1:1 binding model was used to fit the data in GraphPad Prism.

The dissociation rate constant k_off_ of selected dArmRPs was determined by a competition-based FA assay. After establishing a submaximal binding signal (30 nM peptide-GFP fusion, and 50 nM dArmRP to about 80% or the maximum signal, equilibrated for 30 min), synthetic HVR-peptide (KRAS4B 174-188, Thermo Fisher Scientific) is applied in excess (2 µM) as a non-fluorescent competitor, and the decay of anisotropy over time is observed. The competitor was added by the automated injector module of a Tecan Spark reader. Data were fitted with a one-phase exponential decay model to determine k_off_.

### Western blotting

Cells were lysed in RIPA buffer containing 1x Halt protease inhibitor cocktail (Pierce), and 1× Halt phosphatase inhibitor cocktail (Pierce). Cell lysates or recombinant proteins were resolved by SDS-PAGE on Mini-PROTEAN TGX^TM^ 4-20% gels (BioRad) and subsequently transferred onto a nitrocellulose membrane using a Trans-Blot® Turbo^TM^ transfer system (BioRad). Membranes were blocked for 30 min with casein blocking buffer (Sigma-Aldrich). Staining was performed in three steps. First, membranes were stained with HA-tagged dArmRP proteins (5 μg/ml). Further staining was performed with anti-HA primary mouse antibodies (Sigma-Aldrich, #H9658) and rabbit anti-panRAS (Cell Signaling Technology, #3339), and as secondary antibodies, sheep anti-mouse DyLight^TM^ 800 (Rockland Immunochemicals, #610-645-002) and goat anti-rabbit AF® 680 (Invitrogen, #21076). Imaging was performed using an Odyssey CLx instrument (LI-COR Biosciences).

### Enzyme-linked immunosorbent assay (ELISA)

The indicated biotinylated proteins were immobilized indirectly via neutravidin on Nunc MaxiSorp^TM^ flat bottom 96-Well Plate (Thermo Fisher Scientific) and detected in a three-step process. The dArmRP-HA proteins were used as primary detection reagents and were detected via their C-terminal HA-tag with a primary mouse anti-HA antibody (Sigma-Aldrich, #H9658) and a secondary goat-anti-mouse alkaline phosphatase-conjugated antibody. Detection was initiated by the addition of 3 mM para-nitrophenylphosphate (PNPP). Signals were read at 405 nm (PNPP) and 540 nm (reference) using an Infinite M1000 Pro plate reader (Tecan).

### Nucleotide exchange assay

SOS-mediated nucleotide exchange was monitored by a quenching resonance energy transfer (QRET) assay in low volume 384-well plates (Corning) (43). KRAS(1–188) (50 nM), and 0-3 µM of each dArmRP were incubated in assay buffer (20 mM HEPES (pH 7.5), 1 mM MgCl_2_, 10 mM NaCl, 0.01% Triton X-100, and 0.005% γ-globulins). The reaction was initiated after 10 min by adding 10 nM Eu^3+^-GTP, 2.5 µM of the soluble quencher MT2 (both from QRET Technologies) and the catalytic domain of SOS (SOScat, 10 nM), resulting in a final volume of 15 µl. After additional 15 min, time-resolved luminescence (TRL) was monitored using a Tecan Spark 20M instrument with excitation and emission wavelengths of 340 and 620 nm, respectively (800 µs delay and 400 µs decay). Data represent mean ± SD (n=3).

### Loading of RAS with nucleotides

KRAS proteins were purified from *E. coli* in the GDP-bound state. Since the GTP-bound form was preferred for assays, the GDP was replaced with the non-hydrolyzable GTP-analogue 5’-guanylyl imidodiphosphate (GppNHp, Jena Biosciences). For this purpose, KRAS proteins were thawed on ice and diluted (5×) in alkaline phosphatase buffer (40 mM HEPES pH 8, 0.1 mM ZnCl_2_, and 200 mM (NH₄)₂SO₄). Up to 5 mg of KRAS protein was mixed with 3 U/mL of calf intestine phosphatase beads (Sigma Aldrich) and a 20-fold molar excess of GppNHp over KRAS. The protein solution was incubated (RT; 3 h) on a roller shaker before centrifugation (4°C; 20,000 × g; 1 min). The supernatant was transferred to a 0.22 µm centrifugal filter (Sigma Aldrich) and centrifuged (4°C; 20,000 × g; 1 min). The filtered protein solution was supplemented with an additional 20-fold molar excess of GppNHp and a 200-fold molar excess of MgCl_2_. The solution was incubated (RT; 1 h) on a roller shaker before applying the solution to a Zeba^TM^ Spin Desalting Column (7 kDa MWCO, 0.5 mL; Thermo Fisher Scientific) to exchange the buffer to 50 mM HEPES (pH 7.4), 10% (w/v) glycerol, 2 mM TCEP, 125 mM NaCl, 10 mM MgCl_2_. Protein concentrations were determined by measuring at 488 nm for sfGFP-fused proteins with a NanoDrop^TM^ spectrophotometer (Thermo Fisher Scientific).

### Crystallization

To generate a complex, dArmRP proteins were mixed with a 1.2-fold excess of synthetic HVR-peptide (KRAS4B 174-188, Thermo Fisher Scientific) at a total protein concentration of 15 mg/ml and set up for crystallization in sitting-drop vapor-diffusion experiments in 96-well plates. Two different ratios of reservoir:protein solution (1:1 and 2:1) in 300 nl drops were used per well and were incubated against 75 µl reservoir solution at 4°C. Crystals grew within 25 days in 0.2 M NH_4_Cl, 20 %w/v PEG 3350 (PDB: 29IQ) or 30 % w/v PEG 3350, 309 mM NH_4_SO_4_ (PDB: 29IS).

The crystals were mounted in cryo-loops from Hampton Research and flash-cooled in liquid nitrogen with ethylene glycol as a further cryoprotectant. X-ray diffraction data were collected at a wavelength of 1.0 Å on beamline X06SA at the Swiss Light Source, Paul Scherrer Institute (PSI), Villigen, Switzerland equipped with an EIGER 16M detector (Dectris, Baden-Wattwil, Switzerland). The data were processed with XDS, AIMLESS, and autoPROC (44–46). Structures were solved by molecular replacement with a truncated armadillo structure (residues 58-174 of a dArmRP structure similar to PDB-ID: 5AEI) as reference model. Calculation of the electron density and refinement were performed using Refmac (47). For model building and preparation of figures we used Coot and PyMOL (48, 49). Protein:protein interactions were analyzed with the help of PDBePISA and Ligplot software (50, 51).

### Structure prediction

The interaction of dArmRPs with either the peptide sequence of the KRAS4B HVR (171–188) or the full KRAS4B protein (1–188) was modeled using the AlphaFold 3 server (29). To refine the predicted models, energy minimization was performed in the Rosetta force field, by applying the Rosetta relax protocol (52, 53). Predicted structures were visually inspected and the predicted structure with the highest ipTM score was used for further analysis or graphical illustration if not noted differently.

### Identification of homologous sequences

Homologous sequences were identified using NCBI’s protein BLAST server against the non-redundant UniProtKB/SwissProt sequence database (54). Only sequences of human proteins were included. Models, uncultured/environmental sample sequences and non-redundant RefSeq proteins (WP) were excluded. The query sequence was “KKKKKKSKTKCVIM” as derived from KRAS4B (174–188). Default search parameters were used.

### Microinjection and live-cell microscopy

Two days prior to injection, HEK293 cells were cultured in Dulbecco’s Modified Eagle Medium (DMEM) supplemented with heat-inactivated fetal calf serum (FCS), 1% (v/v) penicillin, and 100 µg/mL streptomycin. The cultures were maintained at 37 °C with 5% CO₂ in an imaging dish with a glass bottom (D35-14-1.5-N, Cellvis).

One hour before injection, the medium was replaced with Live-Cell Imaging Solution (LCIS) (A14291DJ, Thermo Fisher Scientific) containing 10% FCS and penicillin/streptomycin. Proteins and plasmids were prepared at a concentration of 25 µM and 5 ng/ug in PBS, respectively.

The microinjection and subsequent live-cell-imaging was performed using a Visitron CSU-W1 microscope similarly as described previously (55). The injection setup included an InjectMan 4 and FemtoJet 4i (Eppendorf) with Femtotip II injection capillaries (Eppendorf). The injection was conducted manually in phase contrast illumination with a constant flow, by piercing the cell membrane and retracting the needle immediately after visible injection. Live-cell imaging was carried out in air with 95% humidity and at 37 °C. Differential interference contrast (DIC) and fluorescence confocal microscopy were performed after illumination or laser excitation at 488 nm, 561 nm, and 640 nm using a PlanApo objective with 100× magnification. Images of injected cell positions and different channels were recorded sequentially every 20 minutes for a duration of 12 hours. Flat-field correction was applied to the recorded images using Matlab R2020b. Flat-field images were acquired using fluorescent dyes at a concentration of 100 mg/mL in water: fluorescein sodium salt (Sigma, #46960 25G F, 488 nm channel), Rose Bengal (Sigma, #198250 5G, 561 nm channel), and Brilliant Blue FCD (Sigma, #80717 100MG, 640 nm channel).

## Author Contributions

Conceptualization, A.P, Y.S and J.N.M; investigation, J.N.M, Y.S, V.L.R, A.H, D.W, K.K and P.M; resources, A.P and K.K; writing—original draft preparation, J.N.M; writing—review and editing A.P, Y.S and J.N.M; visualization, J.N.M.; supervision, A.P; project administration, A.P.; funding acquisition, A.P. All authors have read and agreed to the published version of the manuscript.

## Data Availability Statement

Data supporting the findings of this study are available from the corresponding author on reasonable request.

## Acknowledgments

We thank Dominic Esposito, William Gillette and their team from the Frederick National Laboratory for providing the fully-processed KRAS protein and the SOS1 catalytic domain and David Vukovic (Dept. Biochemistry, University of Zurich) for the expression plasmid for EGFP-KRAS(1–188). The figures 1 and 3A were created with BioRender.com.

## References

1. S. P. Mo, J. M. Coulson, I. A. Prior, RAS variant signalling. Biochem. Soc. Trans. 46, 1325–1332 (2018).

2. M. J. Whitley et al., Comparative analysis of KRAS4a and KRAS4b splice variants reveals distinctive structural and functional properties. Sci. Adv. 10, eadj4137 (2024).

3. P. J. Casey, P. A. Solski, C. J. Der, J. E. Buss, P21ras is modified by a farnesyl isoprenoid. Proc. Natl. Acad. Sci. U. S. A. 86, 8323–8327 (1989).

4. M. C. Seabra, Y. Reiss, P. J. Casey, M. S. Brown, J. L. Goldstein, Protein farnesyltransferase and geranylgeranyltransferase share a common alpha subunit. Cell 65, 429–434 (1991).

5. D. B. Whyte et al., K- and N-Ras are geranylgeranylated in cells treated with farnesyl protein transferase inhibitors. J. Biol. Chem. 272, 14459–14464 (1997).

6. V. L. Boyartchuk, M. N. Ashby, J. Rine, Modulation of Ras and a-factor function by carboxyl-terminal proteolysis. Science 275, 1796–1800 (1997).

7. Q. Dai et al., Mammalian prenylcysteine carboxyl methyltransferase is in the endoplasmic reticulum. J. Biol. Chem. 273, 15030–15034 (1998).

8. S. Dharmaiah et al., Structural basis of recognition of farnesylated and methylated KRAS4b by PDEŒ¥. Proc. Natl. Acad. Sci. U. S. A. 113, E6766–E6775 (2016).

9. M. R. Philips, Ras hitchhikes on PDE6δ. Nat. Cell Biol. 14, 128–129 (2012).

10. A. A. Adjei et al., Phase II study of the farnesyl transferase inhibitor R115777 in patients with advanced non-small-cell lung cancer. J. Clin. Oncol. 21, 1760–1766 (2003).

11. E. S. Kim et al., Phase II study of the farnesyltransferase inhibitor lonafarnib with paclitaxel in patients with taxane-refractory/resistant nonsmall cell lung carcinoma. Cancer 104, 561–569 (2005).

12. E. Van Cutsem et al., Phase III trial of gemcitabine plus tipifarnib compared with gemcitabine plus placebo in advanced pancreatic cancer. J. Clin. Oncol. 22, 1430–1438 (2004).

13. M. Baranyi et al., Farnesyl-transferase inhibitors show synergistic anticancer effects in combination with novel KRAS-G12C inhibitors. Br. J. Cancer 130, 1059–1072 (2024).

14. H. W. Lee et al., A phase II trial of tipifarnib for patients with previously treated, metastatic urothelial carcinoma harboring HRAS mutations. Clin. Cancer Res. 26, 5113–5119 (2020).

15. A. D. Cox, C. J. Der, M. R. Philips, Targeting RAS membrane association: Back to the future for anti-RAS drug discovery? Clin. Cancer Res. 21, 1819–1827 (2015).

16. D. J. Laderach, D. Compagno, Inhibition of galectins in cancer: Biological challenges for their clinical application. Front. Immunol. 13, 1104625 (2022).

17. R. Levy, M. Grafi-Cohen, Z. Kraiem, Y. Kloog, Galectin-3 promotes chronic activation of K-Ras and differentiation block in malignant thyroid carcinomas. Mol. Cancer Ther. 9, 2208–2219 (2010).

18. F. A. Siddiqui et al., PDE6D inhibitors with a New Design principle selectively block K-Ras activity. ACS Omega 5, 832–842 (2020).

19. A. Tomazini, J. M. Shifman, Targeting Ras with protein engineering. Oncotarget 14, 672–687 (2023).

20. C. Madhurantakam, G. Varadamsetty, M. G. Grütter, A. Plückthun, P. R. E. Mittl, Structure-based optimization of designed Armadillo-repeat proteins. Protein Sci. 21, 1015–1028 (2012).

21. J. Z. Zhang et al., De novo design of Ras isoform selective binders. bioRxiv 10.1101/2024.08.29.610300 (2025).

22. C. Reichen, S. Hansen, A. Plückthun, Modular peptide binding: from a comparison of natural binders to designed armadillo repeat proteins. J. Struct. Biol. 185, 147–162 (2014).

23. S. Hansen et al., Curvature of designed armadillo repeat proteins allows modular peptide binding. J. Struct. Biol. 201, 108–117 (2018).

24. S. Hansen et al., Structure and energetic contributions of a designed modular peptide-binding protein with picomolar affinity. J. Am. Chem. Soc. 138, 3526–3532 (2016).

25. Y. Stark et al., Modular binder technology by NGS-aided, high-resolution selection in yeast of designed armadillo modules. Proc. Natl. Acad. Sci. U. S. A. 121, e2318198121 (2024).

26. D. Garcia-Torres, C. A. Fierke, The chaperone SmgGDS-607 has a dual role, both activating and inhibiting farnesylation of small GTPases. J. Biol. Chem. 294, 11793–11804 (2019).

27. H. L. Hartman, K. A. Hicks, C. A. Fierke, Peptide specificity of protein prenyltransferases is determined mainly by reactivity rather than binding affinity. Biochemistry 44, 15314–15324 (2005).

28. A. Kazi et al., Dual farnesyl and geranylgeranyl transferase inhibitor thwarts mutant KRAS-driven patient-derived pancreatic tumors. Clin. Cancer Res. 25, 5984–5996 (2019).

29. J. Abramson et al., Accurate structure prediction of biomolecular interactions with AlphaFold 3. Nature 630, 493–500 (2024).

30. F. Parmeggiani et al., Designed armadillo repeat proteins as general peptide-binding scaffolds: consensus design and computational optimization of the hydrophobic core. J. Mol. Biol. 376, 1282–1304 (2008).

31. E. Michel, S. Cucuzza, P. R. E. Mittl, O. Zerbe, A. Plückthun, Improved repeat protein stability by combined consensus and computational protein design. Biochemistry 62, 318–329 (2023).

32. T. E. Mattox, X. Chen, Y. Y. Maxuitenko, A. B. Keeton, G. A. Piazza, Exploiting RAS nucleotide cycling as a strategy for drugging RAS-driven cancers. Int. J. Mol. Sci. 21, 141 (2019).

33. J. N. Kapp et al., A nucleotide-independent, pan-RAS-targeted DARPin elicits anti-tumor activity in a multimodal manner. Mol. Oncol. 19, 3266–3286 (2025).

34. D. Yang, A. Singh, H. Wu, R. Kroe-Barrett, Comparison of biosensor platforms in the evaluation of high affinity antibody-antigen binding kinetics. Anal. Biochem. 508, 78–96 (2016).

35. R. M. Haley et al., Lipid nanoparticle delivery of small proteins for potent in vivo RAS inhibition. ACS Appl. Mater. Interfaces 15, 21877–21892 (2023).

36. N. Nayerossadat, T. Maedeh, P. A. Ali, Viral and nonviral delivery systems for gene delivery. Adv. Biomed. Res. 1, 27 (2012).

37. K. W. Teng et al., Selective and noncovalent targeting of RAS mutants for inhibition and degradation. Nat. Commun. 12, 2656 (2021).

38. M. Biancucci et al., The bacterial Ras/Rap1 site-specific endopeptidase RRSP cleaves Ras through an atypical mechanism to disrupt Ras-ERK signaling. Sci. Signal. 11, eaat8335 (2018).

39. M. Simon, U. Zangemeister-Wittke, A. Plückthun, Facile double-functionalization of designed ankyrin repeat proteins using click and thiol chemistries. Bioconjug. Chem. 23, 279–286 (2012).

40. G. Chao et al., Isolating and engineering human antibodies using yeast surface display. Nat. Protoc. 1, 755–768 (2006).

41. P. Ernst et al., Rigid fusions of designed helical repeat binding proteins efficiently protect a binding surface from crystal contacts. Sci. Rep. 9, 16162 (2019).

42. M. Bartkiewicz et al., Non-fluorescent mutant of green fluorescent protein sheds light on the mechanism of chromophore formation. FEBS Lett. 592, 1516–1523 (2018).

43. K. Kopra et al., A homogeneous quenching resonance energy transfer assay for the kinetic analysis of the GTPase nucleotide exchange reaction. Anal. Bioanal. Chem. 406, 4147–4156 (2014).

44. P. Evans, Scaling and assessment of data quality. Acta Crystallogr. D Biol. Crystallogr. 62, 72–82 (2006).

45. W. Kabsch, XDS. Acta Crystallogr. D Biol. Crystallogr. 66, 125–132 (2010).

46. C. Vonrhein et al., Data processing and analysis with the autoPROC toolbox. Acta Crystallogr. D Biol. Crystallogr. 67, 293–302 (2011).

47. P. Skubák, G. N. Murshudov, N. S. Pannu, Direct incorporation of experimental phase information in model refinement. Acta Crystallogr. D Biol. Crystallogr. 60, 2196–2201 (2004).

48. P. Emsley, B. Lohkamp, W. G. Scott, K. Cowtan, Features and development of Coot. Acta Crystallogr. D Biol. Crystallogr. 66, 486–501 (2010).

49. W. L. DeLano, The PyMOL Molecular Graphics System. Schrödinger LLC http://www.pymol.org (2002).

50. R. A. Laskowski, M. B. Swindells, LigPlot+: multiple ligand-protein interaction diagrams for drug discovery. J. Chem. Inf. Model. 51, 2778–2786 (2011).

51. E. Krissinel, K. Henrick, “Detection of Protein Assemblies in Crystals” in Lecture Notes in Computer Science, M. R. Berthold, Ed. (Springer-Verlag Berlin Heidelberg, 2005), vol. 3695, pp. 163--174.

52. P. Conway, M. D. Tyka, F. DiMaio, D. E. Konerding, D. Baker, Relaxation of backbone bond geometry improves protein energy landscape modeling. Protein Sci. 23, 47–55 (2014).

53. L. G. Nivón, R. Moretti, D. Baker, A Pareto-optimal refinement method for protein design scaffolds. PLoS One 8, e59004 (2013).

54. S. F. Altschul et al., Gapped BLAST and PSI-BLAST: a new generation of protein database search programs. Nucleic Acids Res. 25, 3389–3402 (1997).

55. D. Vukovic et al., Protein degradation kinetics measured by microinjection and live-cell fluorescence microscopy. Sci. Rep. 14, 27153 (2024).

