## Supplemental Figures and Tables for "Steric shielding of the KRAS4B hypervariable region enables isoform-specific inhibition of prenylation"

**Institutions**

[1] Department of Biochemistry, University of Zurich, Switzerland

[2] Department of Chemistry, University of Turku, Finland

**Corresponding author:**

Prof. Andreas Plückthun
Dept. of Biochemistry
University of Zurich
Winterthurerstr. 190
8057 Zurich

https://plueckthun.bioc.uzh.ch


**This PDF includes:**

Figures S1 to S4
Tables S1 to S5

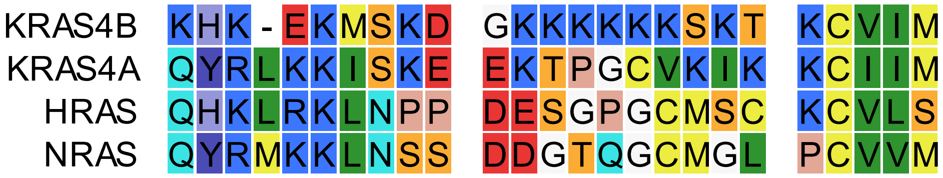


**Figure S1.** Sequence alignment of the hypervariable regions of the four main RAS isoforms.

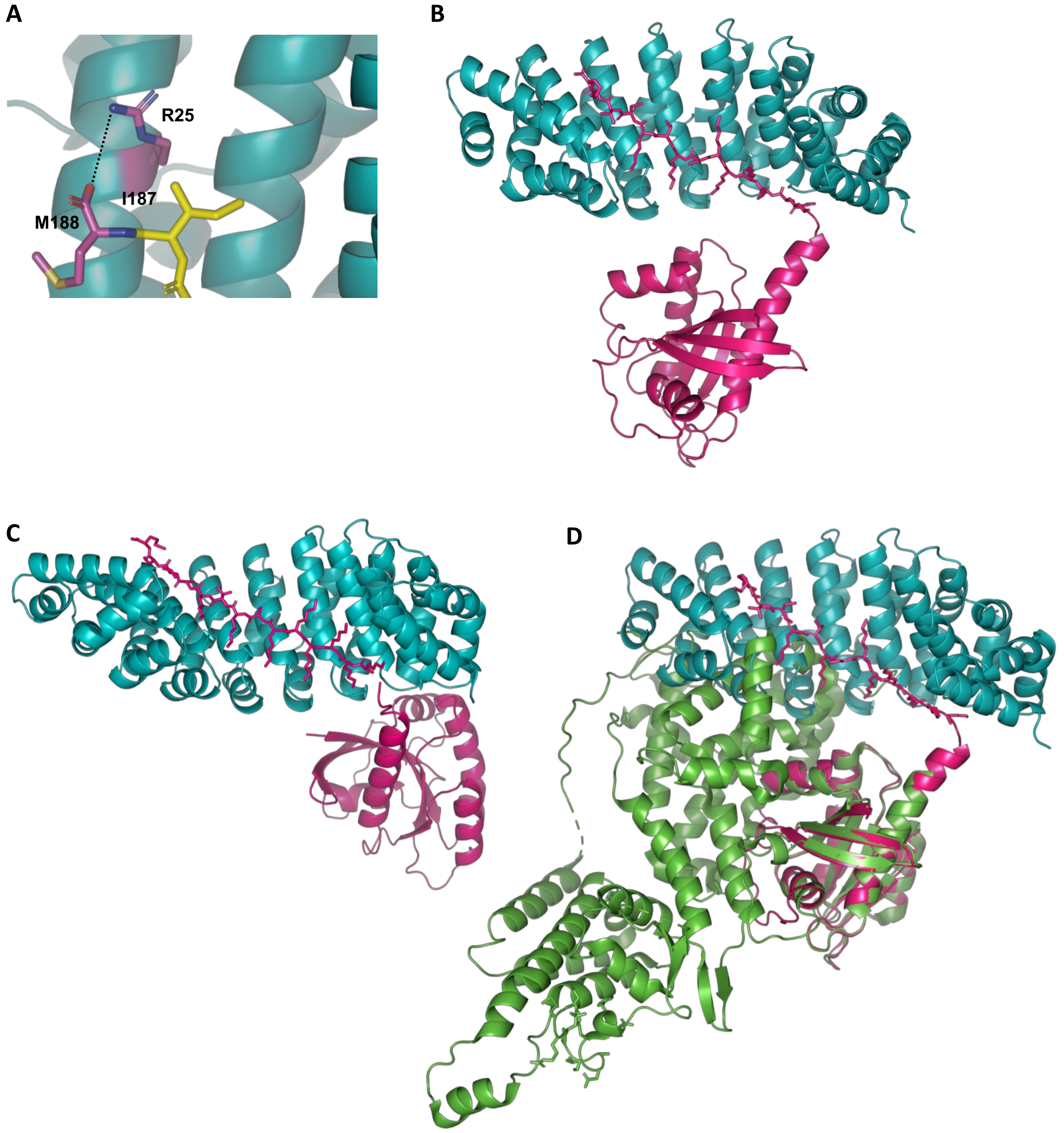


**Figure S2. (A)** The S25R mutation in the N-terminal cap of the selected dArmRPs enables the formation of a salt-bridge with the free carboxyl group of the C-terminal amino acid. **(B)** AF3-generated model of M6_G1 (cyan) bound to KRAS4B (pink). **(C)** AF3-generated model of M7_D5 (cyan) bound to KRAS4B (pink). **(D)** Overlay of SOS1-bound KRAS in green (PDB: 7KFZ) with the predicted structure of the dArmRP M6_G1 (cyan) bound to KRAS4B (pink).

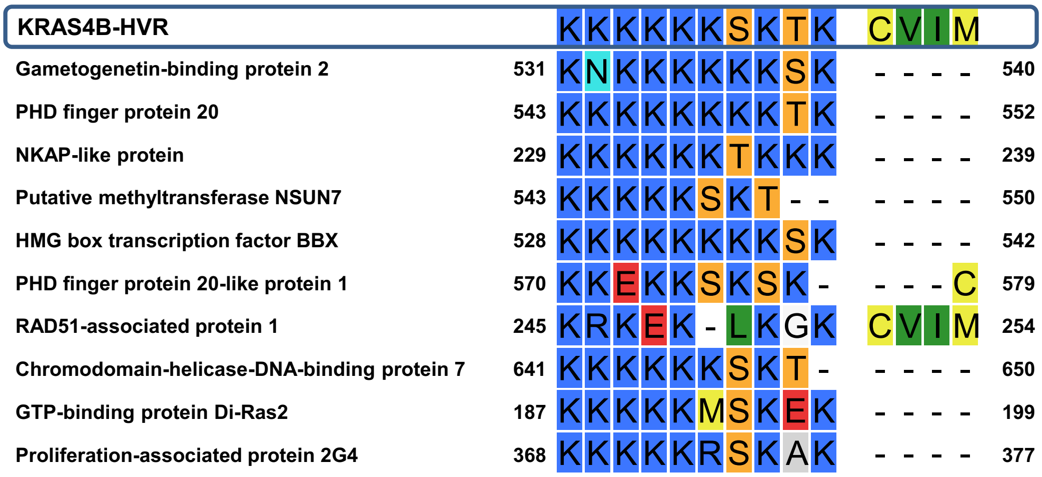


**Figure S3.** Alignment of the ten protein sequences with the highest homology to the residues 175-188 of KRAS4B as identified by protein BLAST.

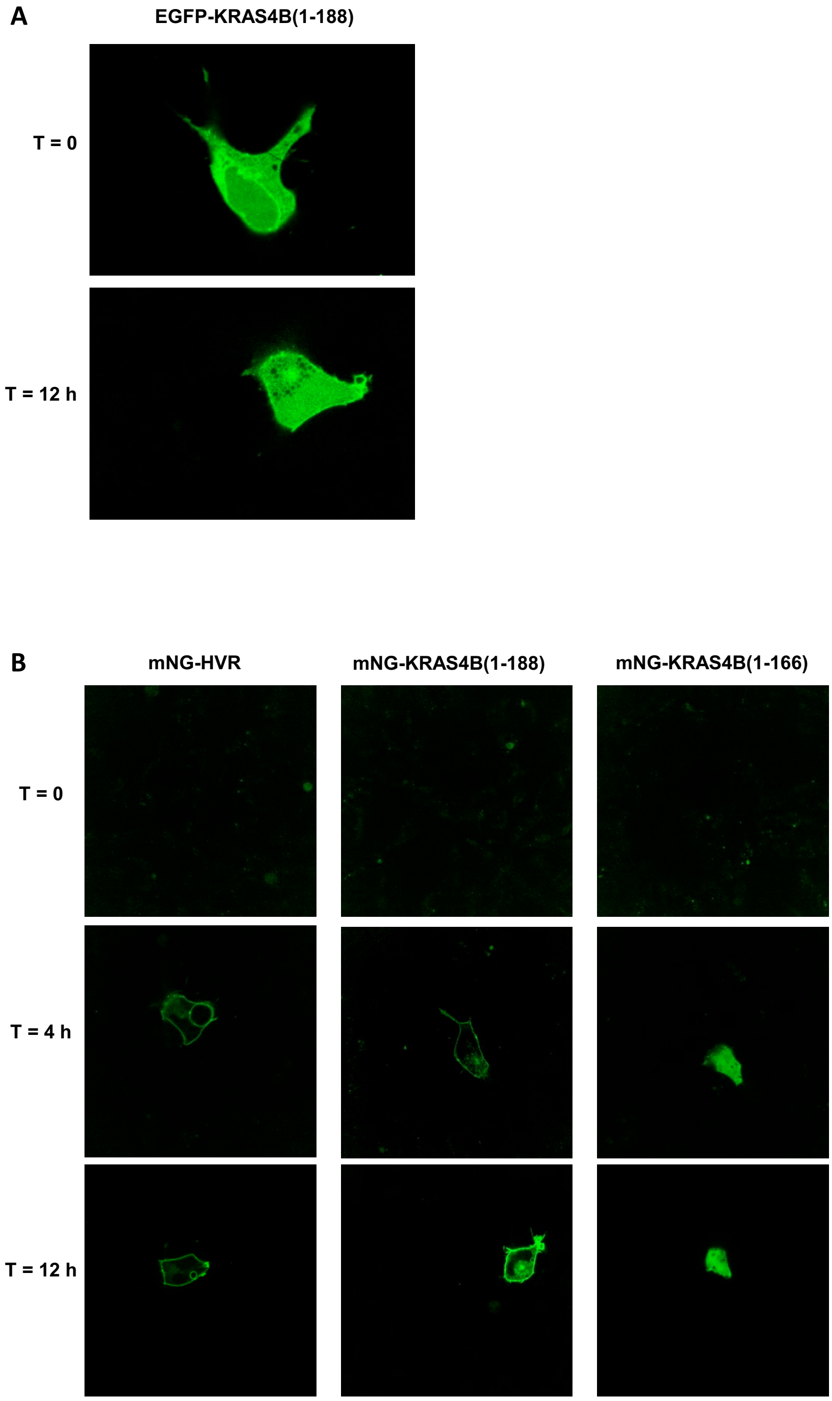


**Figure S4. (A)** Localization of recombinant EGFP-KRAS(1-188) upon microinjection as a function of time. **(B)** Localization of mNeonGreen fusions of the KRAS4V-HVR, KRAS(1-188) and KRAS(1-166) upon expression from a microinjected plasmid as a function of time.

**Table S1.** Selection conditions during affinity maturation.

| | | Concentration during sort | | | | | |
|---|---|---|---|---|---|---|---|
| **Input library** | **Output library** | **EGFP-KRAS4B [nM]** | **KRASless HeLa lysate** | **dGFP [nM]** | **Off-rate selection** | **Sorted Cells** | **Theoretical max. Diversity** |
| parental | G1 F0 | **1st randomization** | | | | | **$9.0 \cdot 10^7$** |
| G1 F0 | G1 F1 | 750 | - | - | no | 923,000 | 923,000 |
| G1 F1 | G1 F2 | 125 | - | - | no | 100,000 | 100,000 |
| G1 F2 | G1 F3 | 125 | yes | - | no | 26,000 | 26,000 |
| G1 F3 | G1 F4 | 100 | yes | 2000 | no | 12,000 | 12,000 |
| G1 F4 | G1 F5 | 62.5 | yes | - | no | 67,000 | 12,000 |
| G1 F5 | G2 F0 | **2nd randomization** | | | | | **$1.4 \cdot 10^6$** |
| G2 F0 | G2 F1 | 62.5 | - | - | no | 163,000 | 163,000 |
| G2 F1 | G2 F2 | 62.5 | - | 625 | no | 72,000 | 72,000 |
| G2 F2 | G2 F3 | 31.25 | yes | 312 | 30 min | 51,000 | 51,000 |

**Table S2.** Affinities of the selected KRAS4B-binding dArmRPs determined by fluorescence anisotropy.

| **dArmRP variant** | **$K_D$ (nM)[a]** | | | |
|---|---|---|---|---|
| | uncleaved, unmodified HVR[a] | cleaved (-VIM), unmodified HVR[b] | cleaved (-VIM), carboxymethylated HVR[c] | cleaved (-VIM), fully modified HVR[d] |
| M6_parental | 874.8 ± 30.0 | 883.2 ± 58.1 | 2234 ± 136 | n.b. |
| M6_G2 | 9.0 ± 1.2 | 136.5 ± 7.8 | 273.9 ± 14.7 | n.b. |
| M6_G1 | 0.4 ± 0.2 | 25.2 ± 1.9 | 36.7 ± 1.5 | n.b. |
| M6_F1 | 0.8 ± 0.2 | 83.9 ± 6.3 | 165.8 ± 7.3 | n.b. |
| M7_A6 | 1.3 ± 0.2 | 35.2 ± 2.4 | 74.8 ± 4.1 | n.b. |
| M7_D5 | 0.5 ± 0.1 | 60.6 ± 3.9 | 147.3 ± 5.4 | n.b. |
| M7_D6 | 0.6 ± 0.3 | 93.2 ± 3.8 | 150.5 ± 5.4 | n.b. |
| M8_F6 | 1.4 ± 0.2 | 32.8 ± 2.5 | 95.0 ± 2.3 | n.b. |
| M8_F8 | 1.3 ± 0.3 | 33.0 ± 1.9 | 77.2 ± 2.4 | n.b. |
| M8_D8 | 0.5 ± 0.1 | 39.4 ± 2.1 | 124.5 ± 3.5 | n.b. |
| M8_E6 | 0.7 ± 0.2 | 36.1 ± 1.8 | 85.9 ± 1.7 | n.b. |

n.b., no binding
[a] N-terminally fused to sfGFP, KRAS4B (165-188)
[a] N-terminally fused to sfGFP, KRAS4B (165-185)
[b] N-terminally fused to 5-FAM, KRAS4B (165-185, Me)
[b] N-terminally fused to 5-FAM, KRAS4B (165-185, FMe)

**Table S3.** X-ray crystallography data collection and refinement statistics

| PDB ID: | M6_G1, trigonal<br>29IQ | M6_G1, tetragonal<br>29IS |
|---|---|---|
| **Data statistics** | | |
| Wavelength | 1.0000 | 1.0000 |
| Resolution | 74.90 - 2.07<br>(2.29-2.07) | 49.48 - 2.84<br>(3.02 - 2.84) |
| Space group | $P3_2$ | $P4_12_12$ |
| Unit cell | 86.497 86.497 206.428<br>90.0 90.0 120.0 | 156.998 156.998 278.995<br>90.0 90.0 90.0 |
| Total reflections | 714674 (29664) | 2244050 (325493) |
| Unique reflections | 79286 (3964) | 78077 (12516) |
| Multiplicity | 9.0 (7.5) | 28.7 (26.0) |
| Completeness (%) | 95.1 (60.7) | 99.8 (99.0) |
| Mean I/sigma(I) | 9.1 (1.7) | 10.9 (0.8) |
| Wilson B-factor | 43.80 | 83.09 |
| R-merge | 0.140 (1.924) | 0.231 (3.532) |
| R-meas | 0.148 (2.065) | 0.240 (3.677) |
| R-pim | 0.049 (0.738) | n.d. |
| CC1/2 | 0.998 (0.415) | 0.999 (0.561) |
| ISa | 21.99 | 30.98 |
| **Refinement** | | |
| Resolution | 50.01 - 2.07<br>(2.10 - 2.07) | 49.48 - 2.84<br>(2.92 - 2.84) |
| Reflections used in refinement | 75316 (350) | 78077 (5642) |
| Reflections used for R-free | 3962 (21) | 4110 (297) |
| R-work | 0.2283 | 0.2200 |
| R-free | 0.3072 | 0.2504 |
| Number of non-hydrogen atoms | 15207 | 19895 |
| macromolecules | 14922 | 19895 |
| ligands | 0 | 0 |
| solvent | 285 | 0 |
| Protein residues | 2022 | 2700 |
| RMS(bonds) | 0.009 | 0.010 |
| RMS(angles) | 1.67 | 1.75 |
| Ramachandran favored (%) | 96.10 | 98.05 |
| Ramachandran allowed (%) | 3.70 | 1.57 |
| Ramachandran outliers (%) | 0.20 | 0.37 |
| Rotamer outliers (%) | 7.80 | 6.70 |
| Clashscore | 10.98 | 4.89 |
| Average B-factor | 57.61 | 113.89 |
| macromolecules | 57.80 | 113.89 |
| solvent | 47.81 | |

**Table S*4*.** Protein sequences of expression constructs used in this study.

| ID | Protein Sequence |
|---|---|
| **EGFP-KRAS4B (1-188)** | GSMVSKGEELFTGVVPILVELDGDVNGHKFSVSGEGEGDATYGKLTLKFICTTGKLPVPWPTLVTTLTYGVQCFSRYPDHMKQHDFFKSAMPEGYVQERTIFFKDDGNYKTRAEVKFEGDTLVNRIELKGIDFKEDGNILGHKLEYNYNSHNVYIMADKQKNGIKVNFKIRHNIEDGSVQLADHYQQNTPIGDGPVLLPDNHYLSTQSALSKDPNEKRDHMVLLEFVTAAGITLGMDELYKGGGGSGMTEYKLVVVGAGGVGKSALTIQLIQNHFVDEYDPTIEDSYRKQVVIDGETCLLDILDTAGQEEYSAMRDQYMRTGEGFLCVFAINNTKSFEDIHHYREQIKRVKDSEDVPMVLVGNKCDLPSRTVDTKQAQDLARSYGIPFIETSAKTRQGVDDAFYTLVREIRKHKEKMSKDGKKKKKKSKTKCVIM |
| **sfGFP-HVR** | GSMSKGEELFTGVVPILVELDGDVNGHKFSVRGEGEGDATNGKLTLKFICTTGKLPVPWPTLVTTLTYGVQCFSRYPDHMKRHDFFKSAMPEGYVQERTISFKDDGTYKTRAEVKFEGDTLVNRIELKGIDFKEDGNILGHKLEYNFNSHNVYITADKQKNGIKANFKIRHNVEDGSVQLADHYQQNTPIGDGPVLLPDNHYLSTQSVLSKDPNEKRDHMVLLEFVTAAGITHGMDELYKGGASKHKEKMSKDGKKKKKKSKTKCVIM |
| **Avi-KRAS (1-166)** | MAGLNDIFEAQKIEWHEGSMGSMTEYKLVVVGAGGVGKSALTIQLIQNHFVDEYDPTIEDSYRKQVVIDGETCLLDILDTAGQEEYSAMRDQYMRTGEGFLCVFAINNTKSFEDIHHYREQIKRVKDSEDVPMVLVGNKCDLPSRTVDTKQAQDLARSYGIPFIETSAKTRQGVDDAFYTLVREIRKLAHHHHHH |
| **Avi-KRAS (1-188)** | GSGLNDIFEAQKIEWHEGGGGSGGGGSGSGMTEYKLVVVGAGGVGKSALTIQLIQNHFVDEYDPTIEDSYRKQVVIDGETCLLDILDTAGQEEYSAMRDQYMRTGEGFLCVFAINNTKSFEDIHHYREQIKRVKDSEDVPMVLVGNKCDLPSRTVDTKQAQDLARSYGIPFIETSAKTRQGVDDAFYTLVREIRKHKEKMSKDGKKKKKKSKTKCVIM |

**Table S5**. EP-PCR parameters for the generation of the libraries G1 F0 and G2 F0.

| **Components** | **Volume (□L)** | **Concentration** |
|---|---|---|
| 10 · Mutazyme II Buffer | 5 | 1· |
| $ddH_2O$ | x | - |
| forward primer (HR1-for) | 2.5 | 0.5 µM |
| reverse primer (HR2-rev) | 2.5 | 0.5 µM |
| template DNA[a] | x | x |
| 40 mM dNTP mix | 1 | 200 µM each |
| Mutazyme II DNA pol. | 1 | 2.5 unit |
| | 50 | |

| **Step** | **Time (s)** | **Temp. (°C)** | **Cycles** |
|---|---|---|---|
| Initial Denaturation | 120 | 95 | 1 |
| Denaturation | 20 | 95 | |
| Annealing | 20 | 56 | 25, 30 |
| Elongation[b] | x | 72 | |
| Final Elongation | 600 | 72 | 1 |
| Hold | ∞ | 8 | 1 |

[a] amount according to library generation (see below)

[b] an elongation time of 60 s per kbp was used and adjusted to the length of the amplified region

| **library name** | **reaction volume** | **amplification cycles** | **template amount** | **mutations per insert** |
|---|---|---|---|---|
| G1 F0 | 50 uL | 25 | 500 ng | 1 - 2 |
| G1 F0 | 50 uL | 30 | 500 ng | |
| G2 F0 | 50 uL | 25 | 50 ng | 4 - 5 |
| G2 F0 | 50 uL | 30 | 50 ng | |
